# Monitoring the battlefield: Antimicrobial resistance, antibiofilm patterns and virulence factors of *Acinetobacter baumannii* isolates from hospital system

**DOI:** 10.1101/2022.12.14.520451

**Authors:** Silas Onyango Awour

## Abstract

Hospital-acquired contagions are on the increase and are a substantial cause of clinical and financial burden for healthcare systems. While contagion control plays a major role in curtailing the spread of outbreak organisms, it is not always effective. One organism of particular concern is *Acinetobacter baumannii*, due to both its persistence in the hospital setting and its ability to acquire antibiotic resistance. *A. baumannii* has appeared as a nosocomial pathogen that displays high levels of resistance to antibiotics, and remains resilient against traditional cleaning measures with resistance to Colistin increasingly reported. Given the costs associated and magnitude with hospital acquired infections, and the rise in multidrug-resistant organisms, it is worth re-evaluating our current approaches and observing for substitutes or aides-de-camp to traditional antibiotics therapies. The aims of this study was to determine antimicrobial resistance, antibiofilm patterns and virulence factors of *Acinetobacter baumannii* isolates from hospital system. Among the five drugs used (Amikacin, Gentamicin, Meropenem, Tigecycline and Tazobactam) it was revealing that all the isolates show resistance to all the drugs. It was revealed that 3 isolates show resistance at difference antimicrobial classes in isolate *05/JOOTRH/22* and *17/JOOTRH/22* were able to produce all type of the virulence enzymes while isolate *11/JOOTRH/22* was able to produce all except protease enzyme. The isolates showed that the biofilm formation inhibitory effects of the various concentrations (0.5, 0.25, 0.125, 0.0625 and 0.03125 mg ml^−1^) were significantly lower than that of the positive control, an indication that biofilm formation was inhibited at these concentrations

## Introduction

Supervision of multidrug-resistant *Acinetobacter spp*. infections is a great task for physicians and clinical microbiologists. Its aptitude to subsist in a hospital setting and its ability to last for extended periods of time on surfaces makes it a frequent cause for health care associated infections and it has led to several outbreaks [1,2]. The organism usually causes varied spectrum of infections that include pneumonia, bacteremia, meningitis, urinary tract infection, and wound infection.

In 1911 in Delft, Nether-lands, a Dutch microbiologist isolated Acinetobacter in Beijerinck [1], but was not definitively recognized until 1971 [2]. Acinetobacter species were originally treatable with antibiotic monotherapy, but high rates of resistance were noted only four years later in 1975 [1]. Over the years resistance rates have amplified and in the eearly 1990s the first reports of carbapenem resistant isolates were documented [3]. Although often still sensitive to colistin, increasingly colistin-resistant isolates have been reported [3,4]. Contamination of mechanical ventilators, hemofiltration machines, cleaning equipment, cleaning fluids, door handles, patient beds, bedside cupboards, and computer keyboards have been reported during outbreaks [5,6]. Strategies exist on explicit actions for detecting and controlling transmission, but despite these guidelines A. baumannii continues to cause nosocomial outbreaks [6], and the emergence of multi-drug resistant (MDR), and extensively-drug resistant (XDR) strains of *A. baumannii* have further raised the stakes for control of this problematic pathogen

During the early 1970s the clinical isolates of Acinetobacter spp. were usually susceptible to gentamicin, minocycline, nalidixic acid, ampicillin, or carbenicillin, singly or in a combination therapy. However, since 1975, increasing resistance started appearing in almost all groups of drugs including the first and second generation cephalosporins. Initially they retained at least partial susceptibility against the third and fourth generation cephalosporins, fluoroquinolones, semi synthetic aminoglycosides, and carbapenems, with almost 100% isolates retaining susceptibility to imipenem. However, during late 1980s and 1990s, worldwide emergence and spread of Acinetobacter strains resistant to imipenem further limited the therapeutic alternatives [7-11]. By the late 1990s, carbapenems were the only useful agents remaining that could combat many severe Acinetobacter infections. Furthermore, due to the emergence of carbapenem resistance in the strains of *A. baumannii*, largely through a clonal spread, the therapeutic options are decreasing [12-14]. Multiple mechanisms have been found to be responsible for the resistance to carbapenems in *A. baumannii*. The mechanisms of antimicrobial resistance in A. baumannii generally falls into three broad categories: [1] antimicrobial-inactivating enzymes, [2] reduced access to bacterial targets (due to decreased outer membrane permeability caused by the loss or reduced expression of porins, overexpression of multidrug efflux pumps) and [3)] mutations that change targets or cellular functions (alterations in penicillin-binding proteins; PBPs) [14, 15]. A combination of several mechanisms may be present in the same microorganism, as has also been observed in other gram-negative bacteria [14].

## Materials and Methods

An examining study was carried out at the Microbiology Laboratory of JOOTRH, Kisumu county-Kenya on the stock isolates which shows Multidrug resistance in the last quarter. The samples were thawed and inoculated on blood agar (BA), Mac-conkey agar (MA) and chocolate agar (CA) using standard protocol [18]. BA and MacConkey agar plates were incubated aerobically, while CBA plates were incubated in anaerobic condition at 37^0^C for 24 to 48 hours in the incubator. The isolates were identified by colony morphology, Gram staining reaction and the biochemical properties [18]. The antimicrobial susceptibility test of isolates was performed by disk diffusion technique method using the standard guidelines and interpretive criteria of the CLSI (2022) [19]. The tests were performed by making a series of antibiotic concentrations on Mueller–Hinton agar plates. A reference strain, *P. aeruginosa spp*. ATCC^®^ 12934 and *S. aureus* ATCC^®^ 29213 were used as control. All the data were entered in SPSS version 20 and Excel 2019. Statistical analyses were done using the same software.

### Biofilm formation inhibition assay

As described by Awuor et al. [16], microtiter plate assay was performed to quantify the effect of commonly used antibiotics on the biofilm formation of *Acinetobacter baumannii*. strains. The test bacteria were first inoculated on Luria-Bertani medium (LB) agar and incubated at 37 °C overnight. Then a colony was identified, picked and inoculated in 10 ml of LB broth and incubated at 37 °C overnight while shaking at 100 r.p.m. for 18 h. By use of a parafilm the flat-bottomed polystyrene tissue culture microplate was sealed for purposes of preventing medium evaporation. After 48 h incubation, the wells were carefully rinsed with double-distilled water to remove loosely attached cells. The microplate was air-dried for 1 h before adding 200 μL per well of 0.4 % crystal violet (CV) solution to the adhered cells in the wells and then stood at room temperature for 15 min. Excess stain was removed by rinsing the wells gently with 200 μL per using distilled water. This was repeated thrice. The microtiter plate was then air-dried for 1 h after, followed by addition of 200 μL of absolute ethanol to each well to solubilize the dye. The OD was measured at OD_590nm_ using a Safire Tecan-F129013 Microplate Reader (Tecan, Crailsheim, Germany). For each experiment, background staining was corrected by subtracting the crystal violet bound to un-treated controls (Blank) from those of the tested sample. The experiments were done in triplicate and average OD_590nm_ values were calculated. To estimate the antibiofilm activity (Abf A) of a given antibiotic the following equation was used

### Detection of other virulence factors of *Acinetobacter baumannii*

#### Detection of haemolysin

Haemolysin production by the *Acinetobacter baumannii isolates* were detected following protocols by Benson et al. [16]. The β-haemolytic activity was tested for on base agar (Himedia, India) supplemented with 7 % sheep erythrocytes for 18–24 h. Pure isolates were cultured on TSA, before streaking on blood agar and further incubated for 24 h at 37 °C. Zones of haemolysis around the colonies indicated the ability of these bacteria to haemolyse RBCs [39].

#### Detection of protease

To detect protease production by the *Acinetobacter baumannii* isolate skim milk agar was used and the protocol that was described in [16]. Briefly, two solutions (A and B) were made and used in this study. Solution A was prepared by adding 10 g skim milk to 90 ml of distilled water then volume was completed to 100 ml gently heated at 50 °C, then autoclaved and cooled to 50–55 °C. And solution B was also prepared by adding 2 g of agar powder to 100 mL of distilled water, mixed thoroughly, then autoclaved and cooled to 50–55 °C. Aseptically, 100 ml of solution A was mixed with 100 mL of solution B. Then the mixture was poured into sterile petri dishes, and then stored at 4 °C until use. This media used to detect the ability of the bacteria to produce protease. The appearance of a cleared hydrolysis zone indicates a positive test [17].

#### Detection of lipase

Lipase production ability by *Acinetobacter baumannii* isolates were determined by methods outlined by Elliot et al. [17]. Briefly, a single colony of an overnight growth was cultured on Rhan medium, and then incubated for 1–5 days at 37 °C. The appearance of a turbid zone around colonies indicates a positive result [16].

#### Detection of lecithinase (phospholipase)

To detect lecithinase, I followed a standard procedure [18]. One pure colony was cultured on a medium of phospholipase activity assay followed by incubation for 1–3 days at 37 °C using established procedures [17]. The appearance of a white to brown colour elongated precipitated zone around colonies is considered a positive result [17].

## Results

The stocked isolates were culture on the Blood Agar media and the growths were observed as shown in the **Plate. 1**, after which a confirmation Identification was done and all the three stock isolates were confirmed to be *Acinetobacter baumannii*. Among the five drugs used (Amikacin, Gentamicin, Meropenem, Tigecycline and Tazobactam) it was revealing that all the isolates show resistance to all the drugs as shows in **Table. 1** and **Plate 2**.

**Plate 1:**
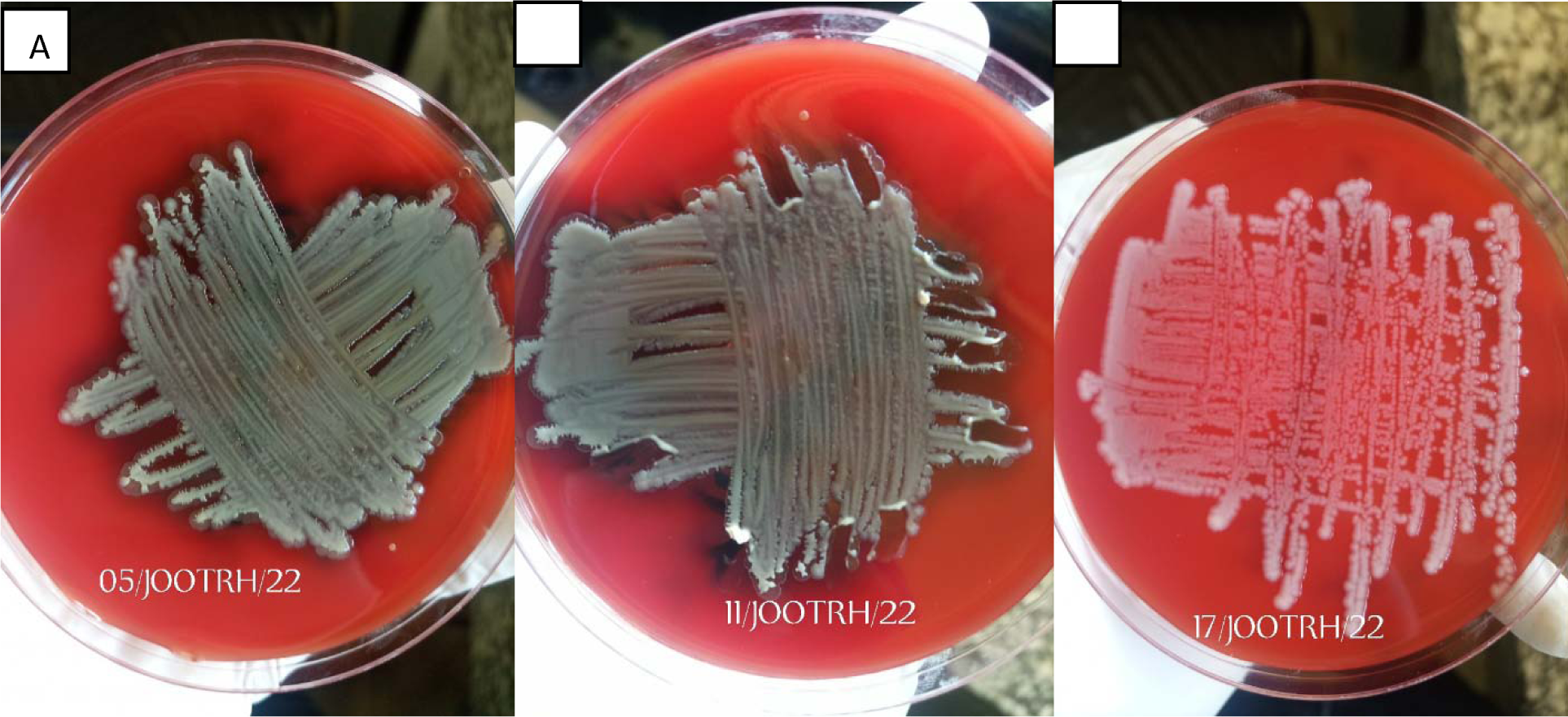
Isolate growth on Blood Agar Media; **A**- *05/JOOTRH/22* growth, **B**- *11/JOOTRH/22* growth, C- *17/JOOTRH/22* growth.

**Table 1:** Antibiotic Resistant Patterns of *Acinetobacter baumannii* Isolates isolated in the study

| *ISOLATES* | *Amikacin* |  | *Gentamicin* |  | *Meropenem* |  | *Tigecycline* |  | *Tazobactam* |  |
| --- | --- | --- | --- | --- | --- | --- | --- | --- | --- | --- |
|  | zone (mm) | interpretation (S, I, R) | zone (mm) | Interpretation (S, I, R) | zone (mm) | Interpretation (S, I, R) | zone (mm) | Interpretation (S, I, R) | zone (mm) | Interpretat (S, I, R) |
| 05/JOOTRH/22 | 6 | R | 11 | R | 6 | R | 10 | R | 6 | R |
| 11/JOOTRH/22 | 8 | R | 10 | R | 6 | R | 11 | R | 6 | R |
| 17/JOOTRH/22 | 11 | R | 6 | R | 6 | R | 12.1 | R | 6 | R |

**Plate 2:**
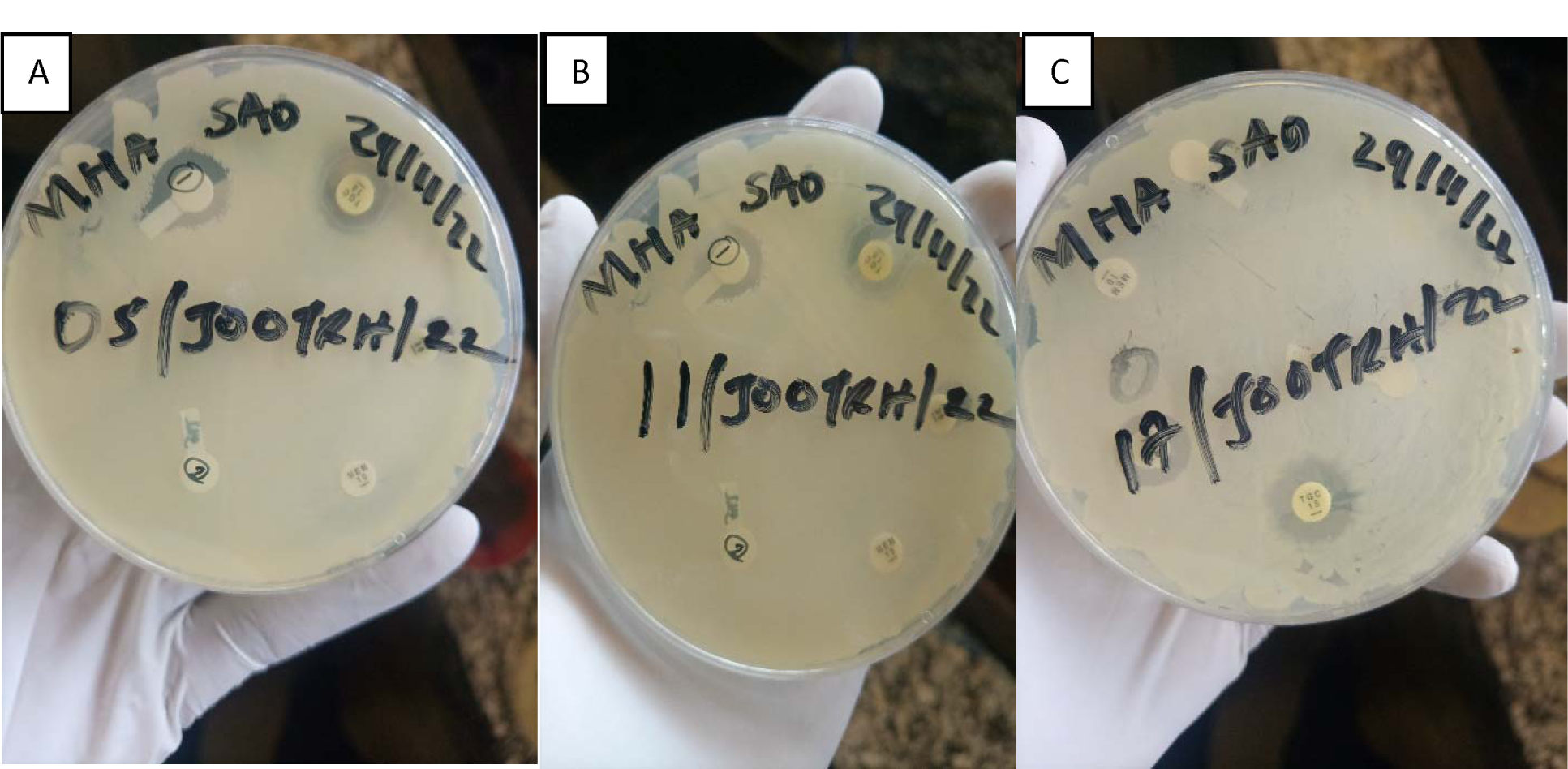
Isolate sensitivity test by use of DDA technique; **A**- *05/JOOTRH/22* sensitivity test, **B**-*11/JOOTRH/22* sensitivity test, **C**- *17/JOOTRH/22* sensitivity test.

This study investigated the production of various virulence enzymes like protease, phospholipase, lipase and haemolysin on the three *Acinetobacter baumannii* isolates which shows MRD and were found to be resistant to common antibiotics used on its management within the study area. It was revealed that 3 isolates show resistance at difference antimicrobial classes in which out of the isolates, isolate **05/JOOTRH/22** and **17/JOOTRH/22** were able to produce all type of the virulence enzymes while isolate **11/JOOTRH/22** was able to produce all except protease enzyme as shown in **Table 2**.

**Table 2:** Detection of some virulence factors of *Acinetobacter baumannii*

| ISOLATE NO. | HAEMOLYSIN | PROTEASE | LIPASE | LECITHANASE |
| --- | --- | --- | --- | --- |
| 05/JOOTRH/22 | + | + | + | + |
| 11/JOOTRH/22 | + | - | + | + |
| 17/JOOTRH/22 | + | + | + | + |

The isolates showed that the biofilm formation inhibitory effects of the various concentrations (0.5, 0.25, 0.125, 0.0625 and 0.03125 mg ml^−1^) were significantly lower than that of the positive control, an indication that biofilm formation was inhibited at these concentrations (**Figs. 1–3**). As much as such inhibitory effects were recorded these findings clearly demonstrate that out of the three isolates that proved to be resistant to commonly used antibiotics, all the isolates have the ability of forming biofilms.

**Fig. 1.**
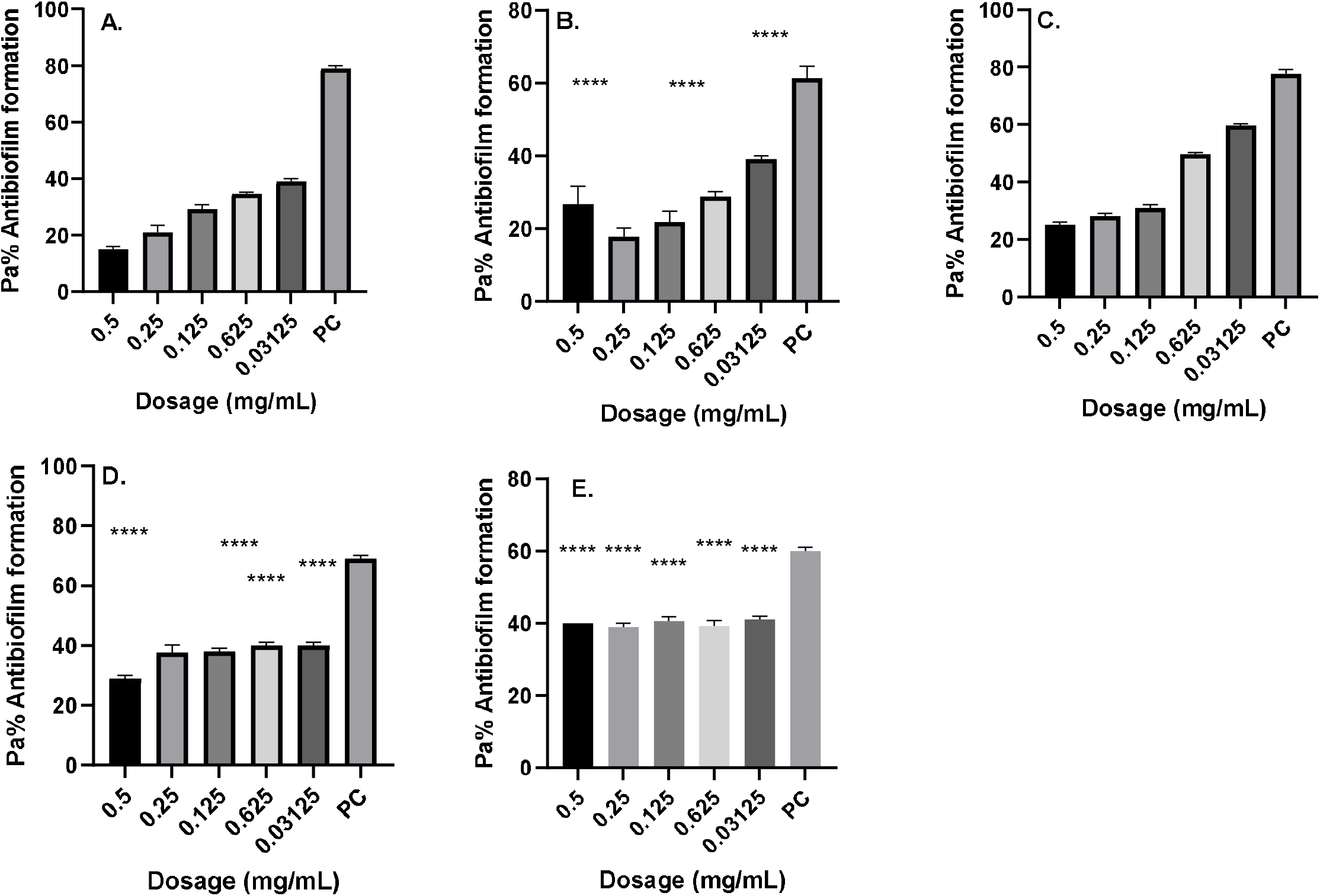
Antibiofilm formation activity against isolate **05/JOOTRH/22** of *Acinetobacter baumannii* against various antibiotics: (a) Gentamicin (b) Amikacin, (c) Tigecycline, (d) Meropenem and (e) Tazobactam; PC=*P. aeruginosa* ATCC^®^ 12934 *–* Positive control (*n*=3, ANOVA Dunnett’s multiple comparisons test; \**P*=0.05; \*\**P*=0.01; \*\*\**P*=0.001; \*\*\*\**P*=0.0001).

**Fig. 2.**
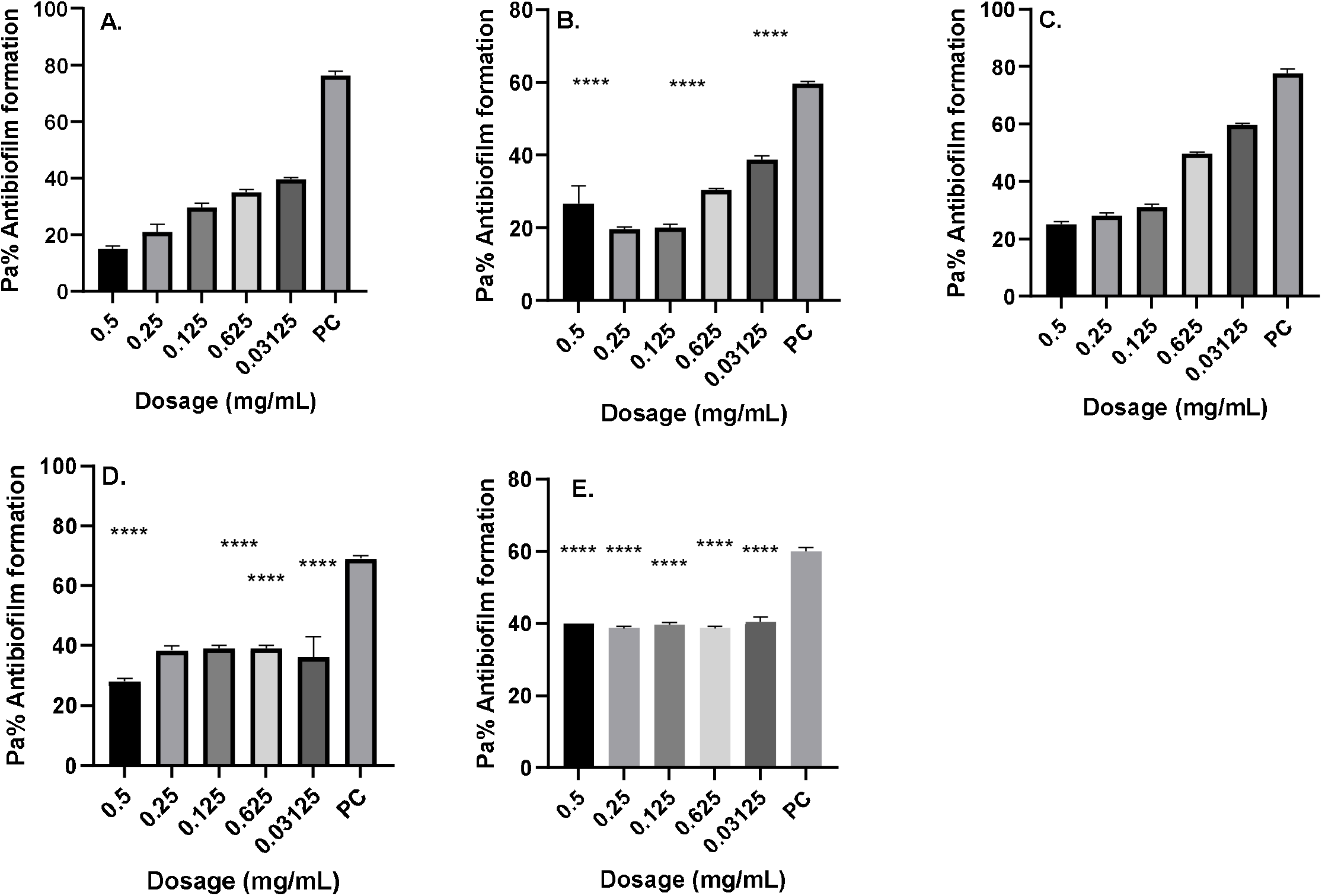
Antibiofilm formation activity against isolate **11/JOOTRH/22** of *Acinetobacter baumannii* against various antibiotics: (a) Gentamicin (b) Amikacin, (c) Tigecycline, (d) Meropenem and (e) Tazobactam; PC=*P. aeruginosa* ATCC^®^ 12934 *–* Positive control (*n*=3, ANOVA Dunnett’s multiple comparisons test; \**P*=0.05; \*\**P*=0.01; \*\*\**P*=0.001; \*\*\*\**P*=0.0001).

**Fig. 3.**
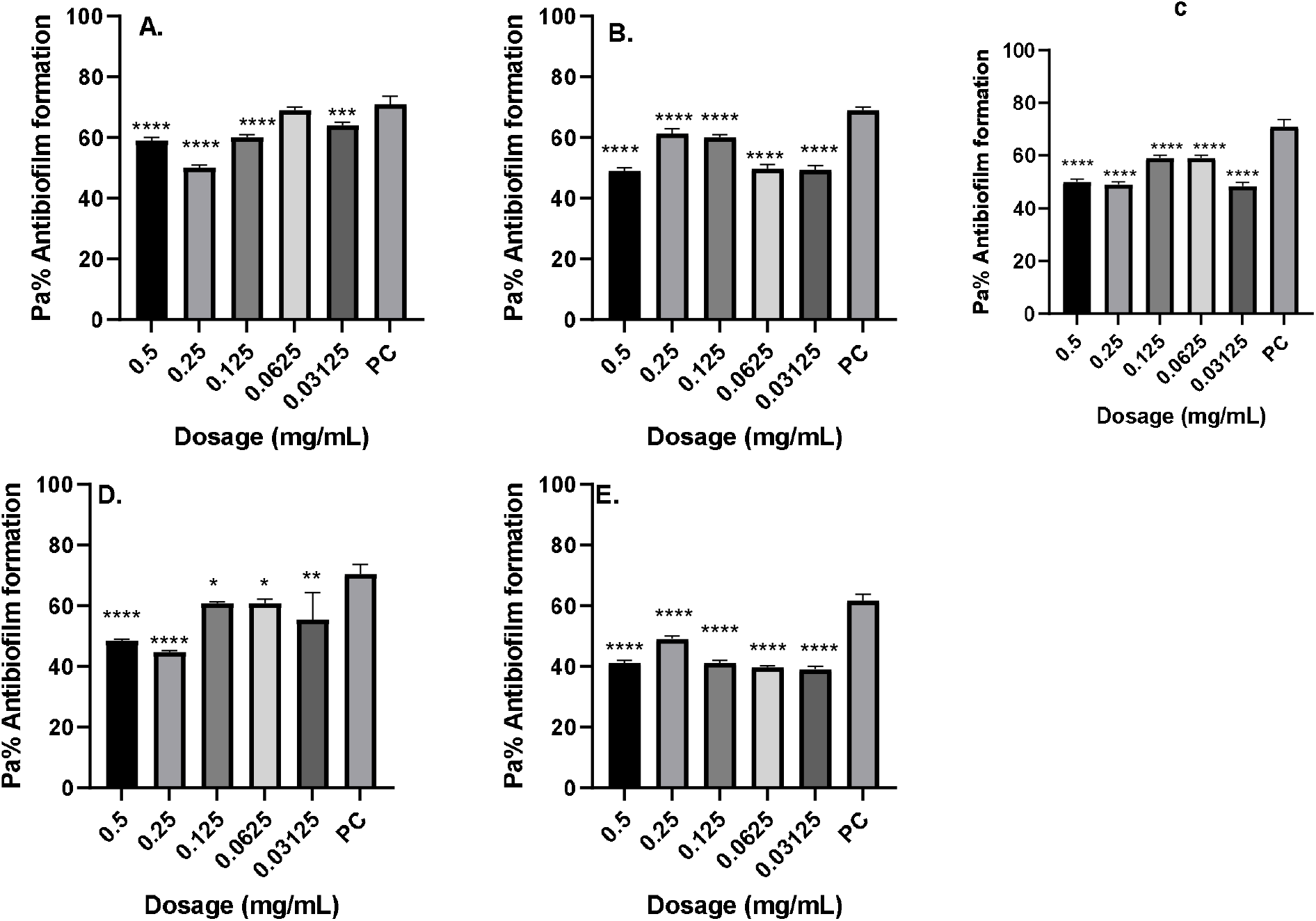
Antibiofilm formation activity against isolate **17/JOOTRH/22** of *Acinetobacter baumannii* against various antibiotics: (a) Gentamicin (b) Amikacin, (c) Tigecycline, (d) Meropenem and (e) Tazobactam; PC=*P. aeruginosa* ATCC^®^ 12934 *–* Positive control (*n*=3, ANOVA Dunnett’s multiple comparisons test; \**P*=0.05; \*\**P*=0.01; \*\*\**P*=0.001; \*\*\*\**P*=0.0001).

Antibiofilm formation activity against isolate **05/JOOTRH/22** of *Acinetobacter baumannii* against various antibiotics like Gentamicin, Amikacin, Tigecycline, Meropenem and Tazobactam was observed in all the antibiotics (Fig. 1). Against Amikacin, Meropenem and Tazobactam a concentration of 0.0125 mg ml^−1^ and 0.0625 yielded significant biofilm formation inhibition (*P*=0.05), while for Meropenem obtained significant differences on the biofilm formation inhibition at a concentration of 0.5 mg ml^−1^ (*P*=0.0001), 0.25 mg ml^−1^ (*P*=0.0001), 0.125 mg ml^−1^ (*P*=0.0001). 0.0625 mg ml^−1^ (*P*=0.0001) and 0.03125 mg ml^−1^ (*P*=0.0001) was observed in all the isolates. It is more worrying that Gentamicin, which is the commonly used antibiotic in the in the hospital, had less inhibitory effects against biofilm formation most so in the isolate **05/JOOTRH/22** and **11/JOOTRH/22**

## Discussion

Carbapenems remain the treatment of choice if isolates retain susceptibility to this antimicrobial class. The MYSTIC surveillance program has documented discordance that favors imipenem as the more potent agent, compared to meropenem, for treatment of MDR Acinetobacter infection and from this study I found resistance to Meropenem and therefore a lot of monitor should be done to Imipenem [20]. Efflux pumps may affect meropenem to a greater degree, whereas, specific beta-lactamases hydrolyze imipenem more efficiently [22].

Tigecycline, a new minocycline derivative, a new glycylcycline agent, received approval from the Food and Drug Administration in June 2005 [21]. The drug is a parenteral, broad-spectrum, bacteriostatic agent and is approved for treatment of complicated skin and skin structure infections as well as intra-abdominal infections caused by susceptible organisms. Tigecycline has activity against the multidrug-resistant Acinetobacter species [22]. From this study I reveal resistance of the drug to the Acinetobacter *baumannii* hence further study should be done on gene determination. Tigecycline’s mechanism of action involves binding to the 30S ribosomal subunit and blocking protein synthesis. High-level resistance to tigecycline has been detected among some MDR Acinetobacter isolates and there is concern that the organism can rapidly evade this antimicrobial agent by upregulating chromosomally mediated efflux pumps [20,15]. Studies have documented overexpression of a multidrug efflux pump in Acinetobacter isolates with decreased susceptibility to tigecycline [20].

Further, the study investigated the ability of these test strains to produce various virulence factors, which may play a role in their pathogenicity. Among the virulence traits examined include detection of proteases, lipases, haemolysin and phospholipase. The study revealed that 100 % of the isolates produced protease, haemolysin and phospholipase while one 1(33.3%) isolates could have not produce lipase. These findings confirm the findings of a previous study [13], which showed that most isolates were protease positive, and that protease enzyme have limited effect on the pathogenesis of this bacteria. Findings from the current study also agree with a study that documented that all isolates had the ability for protease production [23].

Proteases produced by Acinetobacter species have a critical role in pathogenicity, as they are responsible for hydrolysis of several physiologically important proteins such as mucin, fibronectin and lactoferrin [14]. It could also proteolytically activate cholera toxin A subunit, El Tor cytolycin and haemolysin, hence making this pathogen more virulent [15]. A possible explanation to the less activity observed at greater doses in the antibiofilm assay could be associated to the aggregation effects of the antibiotics at site of entry into bacteria cell especially at high dosages something that is not observed at lower dosages. It is likely that aggregation may favor biofilm formation as antibiotics struggle to reach at the point of action and hence bacteria will continue to thrive and hence form more biofilms [24]. This finding agrees with the previous studies on biofilm inhibitions by Taganna et al. [25], which showed higher biofilm inhibitory at lower dosage concentration against the positive control.

## Conclusion

Acinetobacter spp. are rapidly spreading with emergence of extended resistance to even newer antimicrobials. They have the ability to acquire resistance at a much faster pace than other gram-negative organisms. Due to their ease of survival in the hospital environment, they have immense potential to cause nosocomial outbreaks. In addition to antibiotic resistance, their biofilm forming ability plays a crucial role in their in-vitro and in-vivo survival. Thus, to decrease the spread of Acinetobacter infections and reduce the pace of emergence of resistance in MDR Acinetobacter, it is important to promote the rational use of antimicrobials, with implementation and monitoring of the Antibiotics Stewardship Program in hospitals. Hand hygiene and barrier nursing are important to keep the spread of infection in check.

## Supporting information

supplementary doc

