## supplementary doc for "Monitoring the battlefield: Antimicrobial resistance, antibiofilm patterns and virulence factors of *Acinetobacter baumannii* isolates from hospital system"

**SUPPLEMENTARY DATA 1**: JOOTRH results for environmental surface swabs done on 18^th^ /11/22

18 Swabs were collected from critical care, clinical areas and general non-risk areas. Primary culture of the samples was done.

**Results are as follows:**

| **No** | **Description of the swabbing Area** | | **ORGANISM/S ISOLATED** |
| --- | --- | --- | --- |
| 01/JOOTRH/22 | ICU BED 6 | *Acinetobacter baumannii*  *Escherichia coli* | |
| 02/JOOTRH/22 | SINK 2 | *Enterobacter bugandensis*  *Acinetobacter baumannii* | |
| 03/JOOTRH/22 | ICU SLUICE ROOM | *Citrobacter freundii* | |
| 04/JOOTRH/22 | OXYGEN INVENT | No growth obtained | |
| 05/JOOTRH/22 | PATIENT IN ISOLATION BED | *Acinetobacter baumannii* | |
| 06/JOOTRH/22 | WORKING STATION | *Nisseria mucosa*  *Pseudomonas luteola* | |
| 07/JOOTRH/22 | ICU FLOOR | *Bacillus pumilus* | |
| 08/JOOTRH/22 | ICU BED 4 | *Staphylococcus lentus* | |
| 09/JOOTRH/22 | DOOR HANDLES | *Sphingomonas paucimobilis* | |
| 10/JOOTRH/22 | WATER WITH 0.5% JIK | *Staphylococcus caprae* | |
| 11/JOOTRH/22 | SLUICE WASHROOM | *Klebsiella pneumoniae* | |
| 12/JOOTRH/22 | FACE MASK | *Staphylococcus capitis* | |
| 13/JOOTRH/22 | ICU BED 5 | *Staphylococcus lentus* | |
| 14/JOOTRH/22 | OFFICE SWAB | *Staphylococcus haemolyticus* | |
| 15/JOOTRH/22 | ICU VENTILATORS | *No growth obtained* | |
| 16/JOOTRH/22 | ICU OXYGEN MASK | *Micrococcus luteus* | |
| 17/JOOTRH/22 | PATIENT IN ISOLATION BODY FLUID | *Acinetobacter baumannii* | |
| 18/JOOTRH/22 | STAFF CHANGING ROOM | *Staphylococcus epidermidis* | |
